# Mendelian Randomization Analysis Using Mixture Models (MRMix) for Genetic Effect-Size-Distribution Leads to Robust Estimation of Causal Effects

**DOI:** 10.1101/367821

**Authors:** Guanghao Qi, Nilanjan Chatterjee

## Abstract

We propose a novel method for robust estimation of causal effects in two-sample Mendelian randomization analysis using potentially large number of genetic instruments. We consider a “working model” for bi-variate effect-size distribution across pairs of traits in the form of normal-mixtures which assumes existence of a fraction of the genetic markers that are valid instruments, i.e. they have only direct effect on one trait, while other markers can have potentially correlated, direct and indirect effects, or have no effects at all. We show that model motivates a simple method for estimating causal effect (***θ***) through a procedure for maximizing the probability concentration of the residuals, 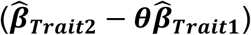, at the “null” component of a two-component normal-mixture model. Simulation studies showed that MRMix provides nearly unbiased or/and substantially more robust estimates of causal effects compared to alternative methods under various scenarios. Further, the studies showed that MRMix is sensitive to direction and can achieve much higher efficiency (up to 3–4 fold) relative to other comparably robust estimators. We applied the proposed methods for conducting MR analysis using largest publicly available datasets across a number of risk-factors and health outcomes. Notable findings included identification of causal effects of genetically determined BMI and ageat-menarche, which have relationship among themselves, on the risk of breast cancer; detrimental effect of HDL on the risk of breast cancer; no causal effect of HDL and triglycerides on the risk of coronary artery disease; a strong detrimental effect of BMI, but no causal effect of years of education, on the risk of major depressive disorder.

## Introduction

Discoveries of genetic susceptibility variants underlying complex traits continue to increase rapidly with ever growing size of genome-wide association studies^1–5^. Mendelian randomization (MR) - a form of instrumental variable analysis for the assessment of the causal effect of one trait on another - provides major opportunity for translation of the increasing knowledge of genetics to improve human health^6,7^. MR analysis has already been widely used for obtaining evidence for drug targets, causal basis for epidemiologic associations and cascading effects among complex molecular traits^8,9^. While MR was originally designed to be used with individual genetic instruments with known biologic functions, the recent trend has been to exploit the multitude of genetic variants emerging from large GWAS. The use of multiple genetic variants can allow to increase the power of MR analysis, correct for weak instrument bias and can provide evidence of causality across a broader set of underlying mechanisms for intervening on the traits^7,8^.

When all of the selected variants satisfy the key assumption of MR analysis, i.e. they only have direct effects on one trait, then the causal effect of that trait on the other can be efficiently estimated by meta-analysis of the well-known ratio estimates of causal effects across the different variants. It is, however, recognized that while availability of many variants provides an opportunity to strengthen MR analysis, there is also major potential for bias as the key assumption, i.e. the variants have no direct effect on the second trait, can be violated due to pleiotropy^7,8^. Indeed, recent empirical studies^3,10–15^ have unequivocally shown that common variants have wide spread pleiotropic effects, and consequently, polygenic MR analysis can be susceptible to bias. Originally, some methods were developed that allow genetic variants to have pleiotropic effects, but they require the strong InSIDE assumption, i.e. the direct and indirect effects are uncorrelated^15–18^. Under this assumption, any genetic correlation between the two traits, as measure by the selected SNPs, can arise solely due to underlying causal relationship between the two traits.

The effects of genetic variants on multiple traits can be correlated when they are mediated through common causal factors, a scenario that is likely to be prevalent in practice. Thus, most recently there has been effort to develop methods that can allow for the presence of invalid instruments which may have complex, possibly correlated pleiotropic effects. In particular, median- and mode-based ratio estimators have been proposed for removing effects of invalid instruments for the estimation of causal effects under different assumptions^19,20^. Further, a recent study proposed the use of methods for outlier detection for conducting sensitivity analysis of ratio estimates to the presence of invalid instruments^21^. While these and other new methods present important progress, there remains important gaps as they can be susceptible to substantial residual bias in the presence of large number of invalid instruments or/and can be inefficient in the regard that they may produce estimates of causal effects with large uncertainty.

In this article, we propose a novel method for estimation of causal effects in multi-marker MR analysis by taking advantage of a working parametric model for the underlying bivariate effect-size distribution of the SNPs across pairs of traits. The model allows genetic correlation to arise from both causal and non-causal relationships, i.e. due to correlated pleiotropic effects. For estimation of causal effects, however, we do not use the likelihood of the data under the model itself to reduce sensitivity to the underlying assumptions. Instead, we propose an estimating equation approach that essentially requires maximization of the probability concentration of the residuals, 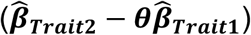, at the “null” component of a normal-mixture model, the form of which is derived from the proposed model. We use extensive simulation studies to show that the proposed method can provide much better trade-off between bias and variance than existing estimators even in a wide set of scenarios. We apply the proposed and existing methods for conducting MR analysis across a variety of exposures and health outcomes using publicly available summary-statistics from very large GWAS. Analysis reveals important differences across methods and new insights to causal relationships underlying some of these traits.

## Methods

### Model setup

We propose a method for two-sample MR analysis that requires only summary-level GWAS association statistics for a putative exposure (Trait 1) and the outcome (Trait 2) of interest from separate studies. We describe the proposed method in the context of independent SNPs. Let 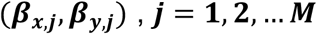, denote the underlying true association coefficients for the ***M*** SNPs for the exposure (***X***) and the outcome (***Y***). The standard MR analysis assumes that all the SNPs are valid instruments, i.e., they are associated with ***X*** but have no direct effect on ***Y***. If the assumption is satisfied, then the two sets of regression coefficients will satisfy a proportional relationship in the form ***β_Y_=θβ_X_***, where ***θ*** is the causal effect of ***X*** on ***Y***. Instead of assuming such a restrictive relationship, we propose modelling the bivariate effect size distribution using a flexible normal-mixture model where the proportional relationship needs to be satisfied only for a fraction of the genetic variants.

We assume a SNP can have four different types of effects: (1) direct effect on ***X*** and an indirect effect on ***Y*** only through ***X***, (2) direct effects on both ***X*** and ***Y***, (3) direct effect on ***Y*** but no relationship with ***X***, (4) related to neither ***X*** nor ***Y*** (**Figure 1**). If we let ***u***_***x***_ and ***u***_***y***_ be the direct effects of a SNP on ***X*** and ***Y***, respectively, then we can write ***β***_***x***_ = ***u***_***x***_ and 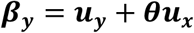. In the above, we note that the second component includes SNPs in “horizontal pleiotropy”, i.e., SNPs with potentially correlated effects on ***X*** and ***Y***. This component allows violation of the InSIDE assumption as we allow ***σ_x,y_* ≠ 0**.

**Figure 1.**
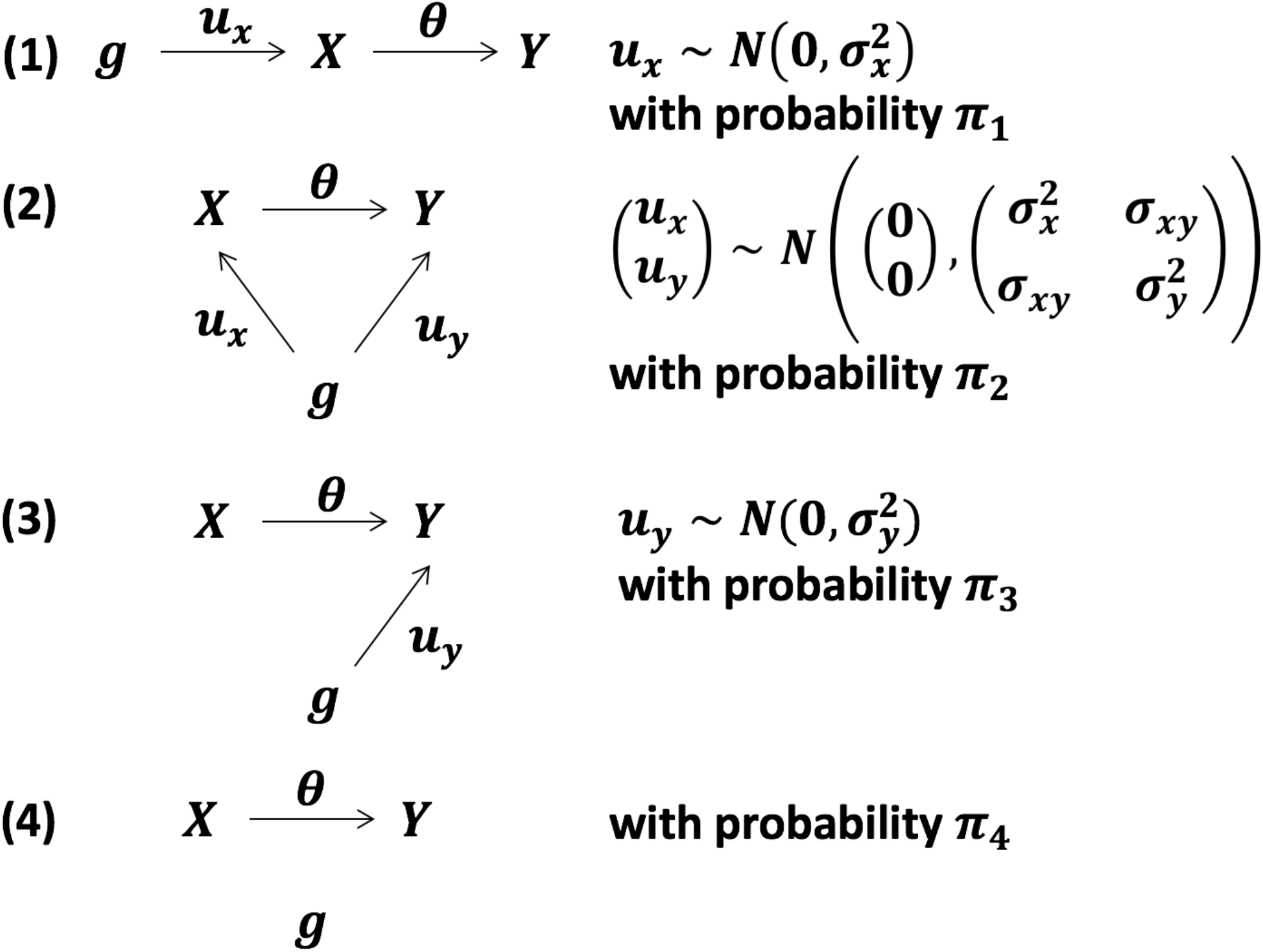
Four components of the Mendelian randomization mixture(MRMix) model. (1) Direct effect on ***X*** and an indirect effect on ***Y*** only through ***X***; (2) direct effects on both ***X*** and ***Y***; (3) direct effect on ***Y*** but no relationship with ***X***; (4) related to neither ***X*** nor ***Y***. Direct effects are denoted by ***u_x_*** and ***u_y_*** and the causal effect is denoted by ***θ***.

### MR analysis using mixture-model (MRMix): a spike detection problem

In GWAS, we obtain noised estimates 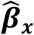 and 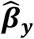, where one can assume 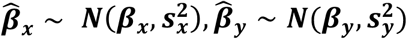, with known standard errors 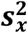 and 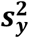 In principle, a likelihood for the observed data can be written by integrating over the “prior” model for the bivariate effect size distribution. However, maximum likelihood estimation of the target parameter θ, jointly with all of the nuisance parameters 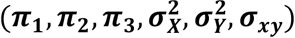 may face computational challenges due to identifiability issues associated with mixture likelihoods. Further, the inference can be sensitive to violation of the underlying modelling assumption.

In the following, we propose an alternative estimation procedure that is computationally simple and rely less on the underlying model assumption. We first observe that that the distribution of the residuals 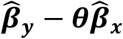 can be written at the true (***θ***) and alternative value (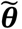)under the proposed model in the form 
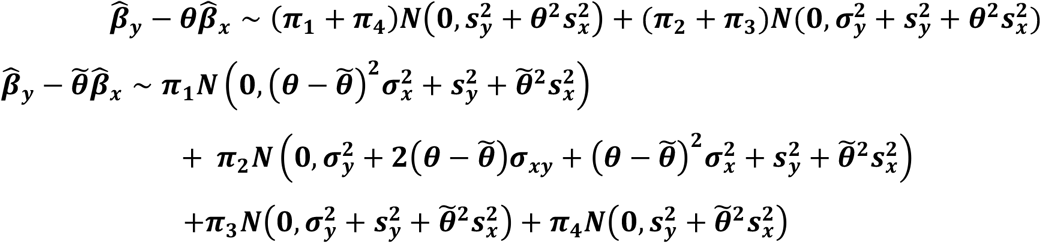

Note that when 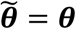, the first and fourth terms collapse, leading to an enrichment of the point mass at 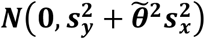. *Note that only at the true value* ***θ***, 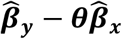 does have an enriched point mass *π*_1_ + *π*_4_ *at* 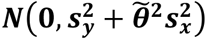, while for other values 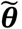 *this point mass is****π_4_***. *The enrichment ***π_1_*** is contributed by the SNPs that have no direct effects on****Y***,*the key assumption underlying instrumental variable (IV) method. Our approach uses this property to identify the causal effect*.

Based on the above observations, we propose the following estimation procedure:

i. For a fixed 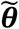 perform maximum-likelihood to fit the two-component normal mixture model in the form 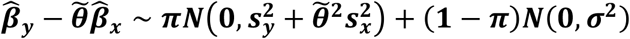 to get estimates of unknown parameters as 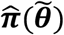 and 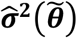;
ii. Search over a grid of 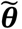 values and choose the one that maximizes 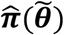 as the estimate, i.e.,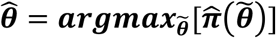.

We note that in step (i), for computational simplification, we are only fitting a two-component normal mixture model which is correct when 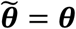, (the true value). When 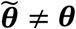, the two-component model is not correct and can only provide an approximation of the underlying multi-component normal mixture model. We observe in a simulation studies that although the model is wrong under the alternative, the proposed estimate 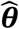 has no asymptotic bias. In contrast, a maximum-likelihood estimator, which maximizes the likelihood of the residuals under the two-component normal-mixture model, produces substantially biased estimate of causal effect due to mis-specification of the model under alternative.

If the study for ***X*** and ***Y*** have overlapping subjects, the null component 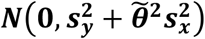 in step (i) can be easily modified to account for correlation in estimated effects. For example, one could use the bi-variate LD score regression^12,22^ to estimate, 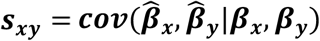, the covariance of GWAS estimates given the effect sizes. Hence the null component could be modified as 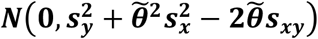 to account for sample overlap across studies of ***X*** and ***Y***.

### Simulation setup

We conducted extensive simulation studies to evaluate the proposed method under different scenarios.

We simulated summary statistics of 200,000 independent SNPs. We first simulate the direct effect sizes (***u_x_***,***u_y_***) and compute the total effect sizes as: 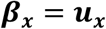 and 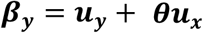. Then we generated the summary statistics by simulating independently from 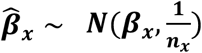 and 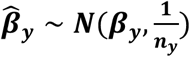, where ***n_x_*** and ***n_y_*** are the sample sizes for studies associated with ***X*** and ***Y***, respectively. This mimics the two-sample MR setup where the exposure and the outcome are measured on independent samples. In all our simulations, we set 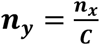 where we vary to ***C = 1,3*** to reflect the fact that for many disease outcomes the effective sample size for ***Y*** can be expected to be lower than continuous exposures ***X***.

We simulated true effect-sizes for the SNPs under the hypothesized four component model as well as more complex models that includes additional mixture components. Under *Scenario A*, we simulated effect sizes ***u_x_*** and ***u_y_*** from the four-component mixture model (Figure 1), where we vary the proportion of valid IVs by changing the
ratio of ***π_1_*** and ***π_2_*** according to the following specifications:

(A.1)50% causal SNPs for ***X*** are valid IVs:***π_1_*** = ***π_2_***
= **0.01.**
(A.2) 25% causal SNPs for ***X*** are valid IVs:***π_1_*** = **0.005**, ***π_2_*** = **0.015.**

For both cases, we set 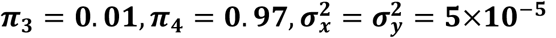 and 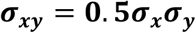. According to the model, a total of 2% of the SNPs have direct effect on ***X*** and either 2% or 2.5% of the SNPs have direct effect on ***Y*** and the overlapping SNPs have strongly correlated effects. The total heritability of ***X***; is 20% and that for ***Y*** ranges from 18.8% to 28.8%, respectively. We allow the causal effect (***θ***) of ***X*** on ***Y*** to be null, or non-null in positive or negative directions so that the genetic correlations due to pleiotropic and causal effects can act in opposite directions.

In *Scenario B*, we simulated effect-sizes using more complex normal mixture model that allows existence of clusters of non-null SNPs with distinctly larger effects than others^4^. In particular, we allow a fraction of causal SNPs for ***X*** to have distinctly larger effects and we assume the SNPs that have larger effects are more likely to be valid IVs. Similarly, among SNPs which have direct effect on ***Y***, we allow existence of SNPs which have distinctly larger effects and these SNPs are less likely to have direct effect on ***X***. We allow SNPs with distinct cluster of effect-sizes through incorporation of additional normal-mixture components with distinct variance-component parameters (see **Supplementary Notes** for more details).

### Inclusion of SNPs with liberal threshold

While in our main analysis, we focused on analysis based on SNPs as “instruments” that have achieved genome-wide significance in the study associated with ***X***, we explored the ability of the method to handle additional SNPs below genome-wide significance. When SNPs are included using more liberal threshold, one would expect a fraction of these SNPs to be “null”. In the presence of null SNPs, the probability concentration of the two-component mixture model for the residuals at the null component is (***π*_1_**+***π*_4_**), where ***π*_1_** is the proportion of valid instruments and ***π*_4_** is the proportion of null SNPs. Thus, while inclusion of more SNPs as potential instruments could lead to increase in efficiency due to increase in the underlying valid instruments, if a very liberal threshold is used, then large value of ***π*_4_** can obscure the enrichment that is expected for the distribution of residual at the null component due to ***π*_1_**. Thus, one would expect there would be is an optimal threshold for SNP selection, as is typically observed for building polygenic risk scores for risk-prediction. We varied the p-value threshold for instrument selection to 0.005, 5×10^-4^, 5×10^-6^ and studied the bias and standard errors for resulting MRMix estimates. Further, as the winner’s curse problem can create bias when selection of SNPs and estimation of their effects are done based on the same study, we also studied the performance of MRMix when effect-sizes of the SNPs associated with ***X*** are estimated based on an independent dataset than the one used to select the SNPs.

### Summary level data

We applied MRMix for the analysis of publicly available GWAS summary level data to explore causal relationships underlying a variety of exposure-outcome pairs of interest. We selected these pairs based on available sample sizes and number of underlying instruments, existing evidence of epidemiologic associations in the literature or/and evidence of causality from recent MR studies. On the exposure side, we accessed data for height and body mass index^5^, blood lipids^23^, education attainment^24^, blood pressure^25^ and age at menarche^26,27^. For data analysis, we only selected SNPs to be potential “instruments” if they reach genome-wide significance (p-value < 5×10^-8^) in the respective studies. Further, we used LD-clumping with an ***r*^2^** threshold of 0.1 to select ae set of independent instruments for each trait. The number of instruments across the different exposures varied between 70–3801, with the largest numbers being available for height (K=3801), BMI (K=973) and blood pressure (K=237) due to availability of results from the large UK Biobank study.

On the outcome side, we accessed data for coronary artery disease (CAD)^28^ for its analysis in relationship to known major risk factors BMI, blood lipids and blood pressure^29^–35; breast cancer in relationship to several known epidemiologic risk-factors, including height, age-at-menarche, BMI, cholesterol level^36^–39; and major depressive respective studies. Further, we used LD-clumping with an disorder (MDD)^40^ in relationship to BMI and years of education^41^–43. In addition, we explored potential causal interrelationship among some of the exposures themselves, such as between BMI and blood pressure, and between BMI and age-at-menarche^44,45^.

For all datasets, we only included SNPs among a set of ∼1.07 million HapMap3 SNPs that have MAF>0.05 and match alleles in the 1000 Genomes European sample. We set the first allele in 1000G data as effect allele, flipping the sign of the z statistic when necessary. We also removed SNPs whose reported sample sizes are less than 2/3 of the 90^th^ percentile of the sample size distribution across SNPs in respective studies to account for imprecision introduced by large amount of missing data. Finally, we remove SNPs in major histocompatibility complex (MHC) region (26∼34Mb on chromosome 6) and SNPs that have very large z score (z^2^ > 80) to prevent the outliers that may unduly influence the results^46^.

### Variance Estimation

At present, in our data analysis we consider a simple bootstrap based variance estimation of the proposed estimator. From the SNPs that pass the p-value screening and LD clumping, we randomly draw 100 bootstrap samples and calculate the MRMix estimates. The variance of the MRMix estimator is calculated as the empirical variance of the 100 estimates from bootstrap samples. Simulation studies show that when the number of available instruments are large, the bootstrap estimator of variance based on re-sampling of SNPs have good accuracy (**Supplementary Table 5**).

### Alternative Methods

For both simulations and real data applications, we compare MRMix with existing popularly used MR methods that allow estimation of causal effects. In particular, we included inverse-variance weighted (IVW) method^47^, weighted median^19^, weighted mode^20^ and Egger regression^16^. Further, we observe that if the InSIDE assumption holds across all SNPs, then the LD score regression methodology^12^ can be used to estimate the causal effect using all SNPs without any pre-selection. In this case, the estimate is simply given by 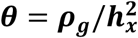, where *ρ_g_* is the estimated genetic covariance of the pair of traits and 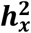 is the estimated heritability of ***X*** (see **Supplementary Notes** for details). Thus, as a benchmark for comparison, in real data analysis, we also report estimate of causal effect based on the LD score regression.

## Results

### Simulations

Simulation studies show that MRMix can be far more robust compared to existing popular methods for MR analysis in a wide range of scenarios (**Table 1, Supplementary Tables 1 and 2**). For example, when genetic correlation due to the causal relationship and pleiotropic effects are in the same direction (**Table 1**), MRMix generally produced nearly unbiased estimates of causal effects as long as the sample size for GWAS for the exposure (***X***) and the corresponding number of instruments reached a minimum threshold (e.g. *n_x_* > 100 *K,K > 100*). The bias was minimal or moderate even when only 25% of the instruments were valid. Among the alternatives, the IVW and the Egger regression methods had the largest bias in directions away from null; the weighted median method was less sensitive, but had considerable bias in many scenarios; and the weighted mode method was the least biased. The bias of the weighted mode method was comparable to that of MRMix in most scenarios when the number of valid instruments were 50%, but were substantially more when the number of valid instruments dropped to 25% and the sample size for the GWAS of the outcome (***Y***) was relatively small (e.g. *n_x_* = 100*K, n_y_*= 33.3 *K*).

**Table 1.**
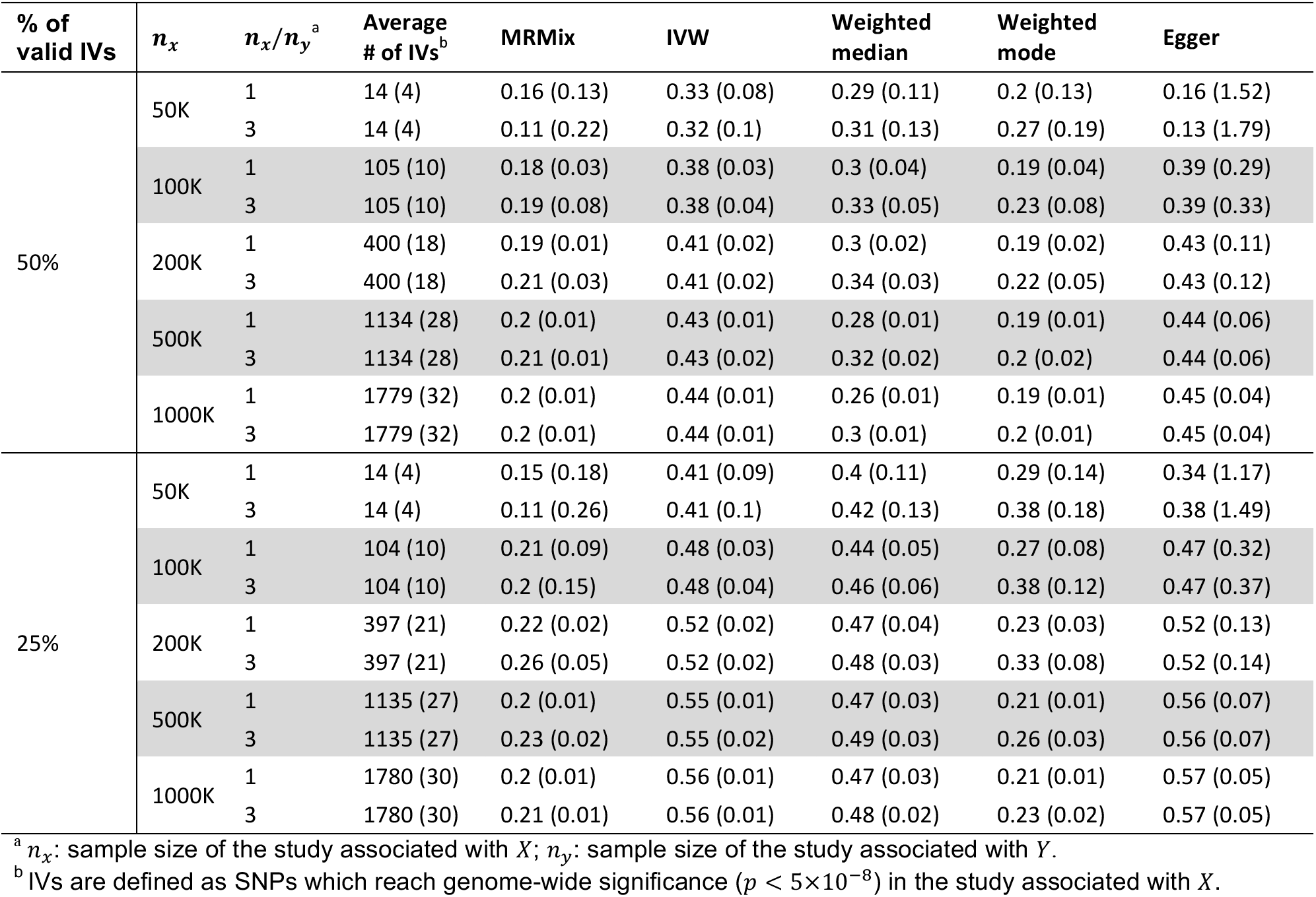
Performance of MRMix and alternative methods for estimation of causal effects (***θ***)in simulation studies. Estimates of association coefficient for SNPs across two traits are simulated assuming an underlying four-component model for effect-size distribution (Scenario A, see **Methods**), where SNPs could have direct effects on neither traits, only on *X*, only on *Y*, or on both with the effects being correlated. The proportion of valid instruments, i.e. the SNPs which have only direct effects on *X*, as a proportion of the total number of SNPs which are associated with *X*, are fixed at 50% or 25%. Mean (standard deviation) of causal estimates are reported over 100 simulations. The true causal effect *θ* is 0.2.

| % of valid IVs | $n_x$ | $n_x/n_y^a$ | Average # of IVs<sup>b</sup> | MRMix | IVW | Weighted median | Weighted mode | Egger |
| --- | --- | --- | --- | --- | --- | --- | --- | --- |
| 50% | 50K | 1 | 14 (4) | 0.16 (0.13) | 0.33 (0.08) | 0.29 (0.11) | 0.2 (0.13) | 0.16 (1.52) |
|  |  | 3 | 14 (4) | 0.11 (0.22) | 0.32 (0.1) | 0.31 (0.13) | 0.27 (0.19) | 0.13 (1.79) |
|  | 100K | 1 | 105 (10) | 0.18 (0.03) | 0.38 (0.03) | 0.3 (0.04) | 0.19 (0.04) | 0.39 (0.29) |
|  |  | 3 | 105 (10) | 0.19 (0.08) | 0.38 (0.04) | 0.33 (0.05) | 0.23 (0.08) | 0.39 (0.33) |
|  | 200K | 1 | 400 (18) | 0.19 (0.01) | 0.41 (0.02) | 0.3 (0.02) | 0.19 (0.02) | 0.43 (0.11) |
|  |  | 3 | 400 (18) | 0.21 (0.03) | 0.41 (0.02) | 0.34 (0.03) | 0.22 (0.05) | 0.43 (0.12) |
|  | 500K | 1 | 1134 (28) | 0.2 (0.01) | 0.43 (0.01) | 0.28 (0.01) | 0.19 (0.01) | 0.44 (0.06) |
|  |  | 3 | 1134 (28) | 0.21 (0.01) | 0.43 (0.02) | 0.32 (0.02) | 0.2 (0.02) | 0.44 (0.06) |
|  | 1000K | 1 | 1779 (32) | 0.2 (0.01) | 0.44 (0.01) | 0.26 (0.01) | 0.19 (0.01) | 0.45 (0.04) |
|  |  | 3 | 1779 (32) | 0.2 (0.01) | 0.44 (0.01) | 0.3 (0.01) | 0.2 (0.01) | 0.45 (0.04) |
| 25% | 50K | 1 | 14 (4) | 0.15 (0.18) | 0.41 (0.09) | 0.4 (0.11) | 0.29 (0.14) | 0.34 (1.17) |
|  |  | 3 | 14 (4) | 0.11 (0.26) | 0.41 (0.1) | 0.42 (0.13) | 0.38 (0.18) | 0.38 (1.49) |
|  | 100K | 1 | 104 (10) | 0.21 (0.09) | 0.48 (0.03) | 0.44 (0.05) | 0.27 (0.08) | 0.47 (0.32) |
|  |  | 3 | 104 (10) | 0.2 (0.15) | 0.48 (0.04) | 0.46 (0.06) | 0.38 (0.12) | 0.47 (0.37) |
|  | 200K | 1 | 397 (21) | 0.22 (0.02) | 0.52 (0.02) | 0.47 (0.04) | 0.23 (0.03) | 0.52 (0.13) |
|  |  | 3 | 397 (21) | 0.26 (0.05) | 0.52 (0.02) | 0.48 (0.03) | 0.33 (0.08) | 0.52 (0.14) |
|  | 500K | 1 | 1135 (27) | 0.2 (0.01) | 0.55 (0.01) | 0.47 (0.03) | 0.21 (0.01) | 0.56 (0.07) |
|  |  | 3 | 1135 (27) | 0.23 (0.02) | 0.55 (0.02) | 0.49 (0.03) | 0.26 (0.03) | 0.56 (0.07) |
|  | 1000K | 1 | 1780 (30) | 0.2 (0.01) | 0.56 (0.01) | 0.47 (0.03) | 0.21 (0.01) | 0.57 (0.05) |
|  |  | 3 | 1780 (30) | 0.21 (0.01) | 0.56 (0.01) | 0.48 (0.02) | 0.23 (0.02) | 0.57 (0.05) |

<sup>a</sup> $n_x$ : sample size of the study associated with $X$ ; $n_y$ : sample size of the study associated with $Y$ .
<sup>b</sup> IVs are defined as SNPs which reach genome-wide significance ( $p < 5 \times 10^{-8}$ ) in the study associated with $X$ .

When the genetic correlations due to causal relationship and pleiotropic effects were in the opposite directions (**Supplementary Table 1**) MRMix showed more notable bias in estimation of causal effect – the direction of bias was generally towards null and did not lead to estimates that were in the opposite direction of the true effect. The degree of bias was more when the number of valid instruments were smaller, but the bias steadily disappeared with increasing sample size irrespective of the proportion of valid instruments. In this scenario, the bias of all the other methods were much more severe and sometimes led to average estimate of causal effects in the opposite direction as that of the true effect. As earlier, the weighted mode method was the most robust among the alternatives considered, and yet it produced substantially more biased estimate of causal effect compared to MRMix in a number of scenarios, especially when the proportion of valid instruments was low.

Simulation studies also reveal MRMix estimates have much higher precision, i.e. smaller standard errors, relative to comparably robust estimators. In particular, the relative efficiency of MRMix compared to the weighted mode estimator, evaluated as the inverse of the ratio of respective variances, reached up to 3–4 fold across the different scenarios considered. This gain in efficiency was larger when higher proportion of the IVs was valid. As expected, the IVW method had the smallest standard errors across all the scenarios, but because it produces severe bias its efficiency is not comparable to that of MRMix. There were also several scenarios where the weighted median estimator had smaller standard error compared to MRMix, but in all of these cases the former method produced substantially more biased estimates. Finally, across all scenarios, the egger regression method produced estimates with much larger standard errors than all the other methods considered.

Reverse directional MR analysis shows that MRMix is highly sensitive to the causal direction (**Supplementary Table 3**). In contrast, the IVW, egger regression and the weighted median method reported estimates of causal effect of substantial magnitude from the outcome (***Y***) to the exposure (***X***) even though no such relationship existed. The weighted mode method, similar to MRMix, was found to be robust in the regard that it estimated the causal effect in the opposite direction close to its expected null value. In alternative simulation scenario, where we allow SNPs with larger effects to be more likely to be valid IVs, the methods rank similarly as described above, although all the methods tend to be less biased (**Supplementary Tables 4**). Simulation studies also showed that when the number of selected instruments were large, the bootstrap method for estimation of standard error through re-sampling of SNPs is generally quite accurate (**Supplementary Table 5**).

### Data Analysis

The MRMix analysis detected significant causal effects of genetically determined LDLC, BMI and blood pressure, but not that for HDL-C and triglycerides, on the risk of coronary artery diseases (CAD) (**Table 2**). There were important differences across the methods in estimates of the causal effect for some of these factors. In particular, both IVW and the weighted median method detected significant causal effects of HDL-C and triglycerides, in directions consistent with known epidemiologic associations. The weighted mode method detected some effect for triglycerides, but the estimate had large standard error and was not statistically significant. The MRMix method estimated the causal effect for both of these lipid factors virtually to be zero. All methods detected causal effect of LDL-C in the expected direction and produced estimate of effect-size in similar range with respect to each other, (OR for CAD per SD unit increase in LDL ranged between 1.3–1.58), but notably lower than those reported by previous MR analysis based on smaller number of genetic instruments^48^. Almost all methods detected causal effect of blood pressure and BMI in directions consistent with epidemiologic studies and produced estimates of effect-size in similar range. Egger regression yielded substantially wider confidence intervals thus leading to statistically non-significant results.

**Table 2.** Estimates and 95% confidence intervals for causal effects of various putative risk-factors on three disease outcomes. Here the effect estimates represent increase in log-OR of the disease per s.d. unit increase in the genetically determined level of the risk-factors.

| Disease | Risk-factors<sup>a</sup> | # of IVs<sup>b</sup> | MRMix | IVW | Weighted median | Weighted mode | Egger | LDSC<sup>c</sup> |
| --- | --- | --- | --- | --- | --- | --- | --- | --- |
| Coronary artery disease | BMI | 973 | 0.37 [0.29, 0.45] | 0.34 [0.3, 0.39] | 0.36 [0.3, 0.42] | 0.25 [-0.06, 0.55] | 0.73 [0.4, 1.06] | 0.44 |
|  | LDL | 153 | 0.32 [0.21, 0.43] | 0.28 [0.21, 0.36] | 0.29 [0.22, 0.35] | 0.25 [0.08, 0.43] | 0.46 [0.1, 0.82] | 0.14 |
|  | HDL | 199 | -0.01 [-0.11, 0.09] | -0.17 [-0.24, -0.1] | -0.07 [-0.14, -0.01] | 0 [-0.14, 0.15] | 0.28 [-0.04, 0.6] | -0.28 |
|  | TG | 127 | -0.02 [-0.14, 0.1] | 0.25 [0.17, 0.32] | 0.2 [0.12, 0.28] | 0.12 [-0.17, 0.41] | 0.28 [-0.08, 0.65] | 0.26 |
|  | SBP | 215 | 0.48 [0.36, 0.6] | 0.44 [0.34, 0.54] | 0.44 [0.35, 0.54] | 0.43 [0.17, 0.69] | 0.5 [-0.18, 1.18] | 0.54 |
|  | DBP | 237 | 0.38 [0.26, 0.5] | 0.4 [0.31, 0.5] | 0.36 [0.28, 0.45] | 0.33 [0.08, 0.59] | 0.3 [-0.37, 0.97] | 0.58 |
| Breast cancer | BMI | 842 | -0.18 [-0.27, -0.09] | -0.1 [-0.16, -0.05] | -0.13 [-0.19, -0.07] | -0.14 [-0.41, 0.13] | -0.2 [-0.57, 0.17] | -0.19 |
|  | Height | 3801 | 0.02 [-0.02, 0.06] | 0.02 [0, 0.04] | 0.02 [0, 0.05] | -0.01 [-0.2, 0.19] | 0.03 [-0.11, 0.18] | 0.08 |
|  | LDL | 126 | 0.07 [-0.12, 0.26] | 0.05 [-0.01, 0.12] | 0.06 [-0.01, 0.12] | 0.07 [-0.09, 0.22] | 0.13 [-0.19, 0.46] | 0.02 |
|  | HDL | 152 | 0.2 [0.11, 0.29] | 0.09 [0.04, 0.15] | 0.11 [0.05, 0.17] | 0.14 [-0.01, 0.29] | -0.04 [-0.28, 0.2] | 0.1 |
|  | TG | 103 | 0.01 [-0.19, 0.2] | -0.07 [-0.14, 0] | -0.08 [-0.16, 0] | -0.11 [-0.28, 0.05] | -0.25 [-0.59, 0.08] | -0.04 |
|  | Age at menarche | 90 | -0.12 [-0.23, -0.01] | 0.01 [-0.08, 0.1] | -0.04 [-0.13, 0.05] | -0.08 [-0.32, 0.16] | 0.08 [-0.55, 0.71] | 0.19 |
| Major depressive disorder | BMI | 972 | 0.35 [0.13, 0.57] | 0.15 [0.1, 0.2] | 0.19 [0.12, 0.25] | 0.35 [0.07, 0.62] | 0.15 [-0.2, 0.5] | 0.14 |
|  | Years of education | 70 | 0.01 [-0.3, 0.31] | -0.21 [-0.37, -0.05] | -0.23 [-0.4, -0.06] | -0.24 [-0.76, 0.28] | -0.73 [-2.17, 0.7] | 0 |

<sup>a</sup> LDL: low-density lipoprotein cholesterol. HDL: high-density lipoprotein cholesterol. TG: triglycerides. DBP: diastolic blood pressure. SBP: systolic blood pressure.
<sup>b</sup> IVs are defined as SNPs which reach genome-wide significance ( $p < 5 \times 10^{-8}$ ) in the study associated with $X$ .
<sup>c</sup> LDSC: LD score regression estimates of causal effects is defined as $\rho_g/h_x^2$ , the ratio between the genetic covariance and the heritability of the exposure (see Supplementary Notes for details).

The MRMix analysis detected significant causal effects of genetically determined BMI, HDL-C, age-at-menarche (AAM), but not those for height, LDL-C and TG, on the risk of breast cancer (BC). There were, again, important qualitative and qualitative differences across methods. MRMix inferred negative causal relationship between increased level of BMI and risk of BC, inconsistent with observed epidemiologic association in the opposite direction. A previous MR analysis^49^ that used fewer genetic instruments had also detected the negative direction of the causal effect, but they reported the estimated effect-sizes to be somewhat stronger (OR for per SD unit increase in BMI reported to be in the range 0.56–0.75 compared to 0.84 by MRMix in the current study). In the current analysis, the estimates of causal effect for BMI on the risk of BC from all the other methods were also in the negative direction, but those obtained from the weighted mode and Egger regression methods did not reach statistical significance due to large confidence intervals.

MRMix method identified increased level of HDL-C to be causally related to higher risk of BC (OR=1.22 per SD increase in HDL-C level). The IVW and weighted median methods also detected these effects in the same direction, but the estimated effectsizes were notably (by 50%) smaller. The weighted mode and Egger regression methods did not detect the effect to be statistically significant due to large confidence intervals. The MRMix was the only method which detected significant causal effect of AAM on the risk of BC and the direction of effect was consistent with known epidemiologic association. Intriguingly, one previous study also noted that the standard IVW analysis does not detect any causal effect of AAM on the risk of breast cancer^50^.

However, significant causal effects, in the same direction as the MRMix, have been reported in previous MR analyses which had adjusted for genetic relationship between AAM with BMI^27^. Finally, we found that none of the methods detected a significant causal relationship between height and risk of BC although epidemiologic studies have consistently reported a positive association.

The MRMix provided important insight into the causal relationship underlying known epidemiologic associations of major depressive disorder (MDD) with BMI and years of education (EDY). The method detected that genetically determined BMI increases the risk of MDD. All the other methods detected the same directional effect, but the magnitude of effect-sizes were notably smaller for the IVW, weighted median and Egger regression compared to the weighted mode and the MRMix method. In contrast to BMI, for genetically determined EDY, the MRMix estimated the causal effect of it on MDD to be virtually null. In contrast, all the other methods except Egger regression estimated reduction of risk of MDD by a factor of 0.8 per SD unit increase in genetically determined EDY, with the estimates from IVW and weighted median method being statistically significant. The Egger regression method provided very large estimate with wide confidence interval. These analyses reveal that MRMix provided qualitatively and quantitatively distinct estimates of causal effects of BMI and EDY on the risk MDD compared to those reported in a recent study that used standard IVW method^40^.

## Discussion

In this article, we develop a novel and powerful method for conducting MR analysis using large number of genetic instruments based on normal-mixture models for effectsize distribution where distinct mixture components are incorporated to allow genetic correlations to arise both from causal and non-causal relationships. To gain robustness against possible model misspecification, we do not directly rely on the likelihood for model-based inference. Instead, we develop an estimating equation approach, that, in essence, involves estimation of causal effect through maximization of the probability concentration of “residuals” - defined by the total effect of SNPs on one trait after subtracting off indirect effects through the other trait - at the “null” component of a two-component normal mixture model. Both simulation studies and extensive data analyses show the method is not only robust, i.e. immune to bias in the presence of large number of invalid instruments, but also highly efficient, i.e. it produces substantially more precise estimates of putative causal effects compared to alternative robust methods. The investigations also show the method is sensitive to the direction of causality and hence suitable for bi-directional MR analysis.

Simulation studies clearly demonstrates superior performance of MRMix compared to a number of existing popularly used methods for MR analysis (**Table 1, Supplementary Tables 1-4**). Stability of the method does require an adequately large sample size for the GWAS of the putative “exposure” of interest so that number of instrument available for analysis is reasonably large (e.g. > 50). Once such threshold is exceeded, the method is highly adaptive in dealing with invalid instruments and can maintain excellent trade-off between bias and efficiency compared to other methods. Even in the presence of large number of invalid instruments, the method often produces unbiased estimates of causal effect and, in settings, where there was notable bias, the bias was generally towards null and disappeared with increasing sample size. In the same settings, the alternative methods generally produce much larger bias, sometimes in the directions away from null and the bias always does not diminish with sample size. Among the alternatives considered, the mode-based ratio estimator shows similar level of robustness as MRMix for large sample size, but yet for smaller sample size, in several settings, it produces distinctly larger bias compared to MRMix. Further, MRMix clearly produces estimates with much smaller standard errors and this gain in efficiency is more pronounced when the number of valid instruments is larger, demonstrating the ability of the method to more effectively use the valid instruments compared to the weighted mode estimator.

The MR analysis of casual relationship of age-at-menarche and the risk of breast cancer provides an important empirical illustration of the strength of MRMix. It has been previously observed that genetic correlation between AAM and BC due to underlying direct causal effect and that due to confounding/mediating effect of BMI acts in opposite directions and thus polygenic MR analysis using standard IVW method could fail to identify the causal relationship^50^. However, when the IVW estimator is adjusted for the relationship of AAM associated SNPs with BMI, evidence of casual effect of inverse relationship between AAM and BC risk, consistent with epidemiologic observation, have been seen^27^. Consistent with previous studies, in our analysis, the standard IVW method produced estimate of causal effect for AAM to be virtual null. The MRMix, although did not explicitly account for BMI, produced estimate of causal effect that is both qualitatively and quantitative similar as those reported from BMI adjusted IVW in previous studies^27^,^50^. The weighted mode method, while pulled the estimate more towards the right direction compared to IVW, the estimate was attenuated and had large standard error resulting in statistically non-significant finding. Further, bi-directional MR analysis between BMI and AAM using MRMix suggested that genetically predicted BMI has an inverse causal effect on AAM, and not the other way (**Supplementary Tables 6 and 7**), and thus it appears that the SNPs which are associated with AAM through BMI, which, on its own, influences the risk of BC, are the underlying invalid instruments. The example demonstrates that MRMix has the ability to produce robust and efficient estimate of causal effects in the presence of potentially unobservable confounding factors.

Additional data analyses also illustrated the distinct property of MRMix compared to alternatives. In particular, the MRMix method estimated the causal effect of HDL and triglycerides on the risk of CHD to be virtually null, while standard IVW and weighted median methods detected these effects to be significantly away from null in directions consistent with known epidemiologic associations. A recent study reported significant putative causal effect of years of education on reducing risk of major depressive disorder based on standard IVW analyis^40^. The MRMix analysis indicated that such a causal effect is virtually non-existent and relationship reported previously likely to have been driven by pleiotropic effects. In contrast, MRMix found the causal effect of genetically determined BMI on increasing the risk of MDD to be notably stronger than that is indicated by the traditional IVW method. Thus, it is likely that that there are common genetic pathways underlying these traits which leads to genetic correlation in opposite directions than that is due to the direct causal effect of BMI on MDD.

Recently a number of alternative methods have been proposed for conducting robust MR analysis in the presence of invalid instruments. One such method, termed as MRPRESSO^15^, applies outlier detection tests to each individual genetic variant and removes potentially invalid instruments. While the method was shown to be highly useful for the detection of bias in reported estimate of causal effects in existing MR analysis, the method can only partially correct for bias and relies on the InSIDE assumption. Further, because the method requires conducting a series of tests and evaluating their significance based on simulations, implementation of it can be time consuming and estimation of uncertainty associated with the final estimator accounting for variation associated with all the different stages can be challenging. Another method proposes to obtain IVW estimators for all possible subsets of genetic instruments and then combine them with a model averaging method with lower weight given to more heterogeneous subsets^51^. While the method is shown to be highly robust as well as powerful, it is currently not scalable for the analysis of large number of instruments which is the focus of current study.

Another study proposed analysis of causal relationship between traits based on genetic relationship, but using a different framework than that for the standard MR analysis. The study defined one trait to be partially or fully causal for another, if there is an underlying genetically determined latent variable which influences both traits, but has a stronger relationship with the first than the second^52^. The study defined moment equations for the estimation of parameters quantifying degree of partial causality using GWAS summary-statistics. We believe this novel framework and more traditional MR hypothesis can complement each other to provide improved understanding of the nature of genetic correlation across traits. The use of latent variable framework, for example, detected evidence of partial causality of several cholesterol traits on blood pressure level. The MRMix analysis, as well as none of the other MR analysis methods, detected any evidence of direct causal effects underlying these traits (see **Supplementary Table 6**). Thus the evidence of partial causality is likely to have been primarily driven by the existence of underlying common genetic pathways which are more strongly related to cholesterol level than blood pressure.

The MRMix method has limitations as well. First, the method relies on certain model for underlying effect-size distribution. As we have noted before, the mis-specification of effect-size distribution is not as critical as we do not use the model directly to perform maximum likelihood estimation, but instead use the model as an efficient way of identifying certain “mode”-based estimator based on underlying estimating equations. Simulation studies show that even when the underlying effect-size distribution has more complexity than the assumed model, the estimation of causal effect parameters can remain relatively robust. Nevertheless, more extensive simulation and theoretical studies are needed to further understand the property of the method under complex but realistic models for effect-size distributions as has been evidenced from recent study^4^. Second, the method does require pre-selection of SNPs as genetic instruments based on p-value for significance of their association. We have observed that as long as the significance threshold is stringent (e.g. p-value < 5×10^-8^), there was not substantial winner’s curse bias due to selection of SNPs and estimation of their coefficients from the same study (**Supplementary Table 8**). While the method, in principle, can be extended to include SNPs with more liberal threshold, it can suffer from winner’s curse bias unless the SNP selection and coefficient estimation are performed based on independent studies (**Supplementary Table 8**). Third, in our current study, we have focused on the analysis of independent SNPs selected from GWAS through stringent LD-pruning after prioritizing by p-values. As we perform MR analysis based on the marginal effects of the individual SNPs, some of which may tag multiple underlying causal SNPs, underlying pattern of LD may cause some bias in MRMix as well the other methods. Further studies are needed to investigate the effect of LD in MR analysis, especially when large number of genetic instruments are used. Further studies are also merited to explore the property of MRMix under more complex causal structure in the data, such as partial causality^52^ and multiple components of causality^51^.

In conclusion, MRMix provides a novel tool for conducting robust and powerful MR analysis using large number of genetic instruments that are now rapidly becoming available from recent expansion of GWAS. We demonstrate through simulation studies, as well as variety of real data analyses, that the method has unique ability to trade-off bias and efficiency for estimation of causal effects in the presence of invalid instruments. Application of MRMix for future MR studies will lead to improved understanding of causal basis of genetic correlation across traits.

